# gMCStool: automated network-based tool to search for metabolic vulnerabilities in cancer

**DOI:** 10.1101/2022.11.03.514827

**Authors:** Luis V. Valcárcel, Edurne San José-Enériz, Raquel Ordoñez, Iñigo Apaolaza, Ana Valcárcel, Leire Garate, Jesús San Miguel, Antonio Pineda-Lucena, Xabier Agirre, Felipe Prósper, Francisco J. Planes

## Abstract

The development of computational tools for the systematic prediction of metabolic vulnerabilities of cancer cells constitutes a central question in systems biology. Here, we present *gMCStool*, a freely accessible and online tool that allows us to carry out this task in a simple, efficient and intuitive environment. *gMCStool* exploits the concept of genetic Minimal Cut Sets (gMCSs), a theoretical approach to synthetic lethality based on genome-scale metabolic networks, including a unique database of thousands of synthetic lethals computed from Human1, the most recent metabolic reconstruction of human cells. Based on RNA-seq data, *gMCStool* extends and improves our previously developed algorithms to predict, visualize and analyze metabolic essential genes in cancer, demonstrating a superior performance than competing algorithms in both accuracy and computational performance. A detailed illustration of *gMCStool* is presented for multiple myeloma (MM), an incurable hematological malignancy. gMCStool could identify a synthetic lethal that explains the dependency on CTP Synthase 1 (CTPS1) in a sub-group of MM patients. We provide *in vitro* experimental evidence that supports this hypothesis, which opens a new research area to treat MM.

## INTRODUCTION

With the increasing coverage and accuracy of reference human genome-scale metabolic networks^1,2^, the development of Constraint-based Modeling (CBM) approaches for different biomedical questions has significantly grown in the last years. One of the central topics in CBM has been cancer metabolism^2–4^, as it constitutes an attractive strategy to gain insights into the underlying metabolic dependencies of tumor cells and systematically predicts vulnerabilities. We can find a plethora of methods in the literature^5^ that first construct context-specific metabolic models (CS-models), based on cancer -omics data, and subsequently, computationally predict gene knockout perturbations that sufficiently decreases growth rate or disrupts a key metabolic task for cellular viability (gene essentiality analysis)^6,7^. These methods have been successfully applied to identify cancer-specific essential genes in different tumors; however, there is still substantial room for improvement, as recently shown in Robinson *et al*., 2020^1^.

In this direction, we released a conceptually different approach based on the concept of genetic Minimal Cut Sets (gMCSs), which does not require the construction of CS-models and more generally exploits the concept of synthetic lethality^8,9^. In particular, gMCSs define minimal set of genes whose knockout would render the functioning of a given metabolic task impossible. When they are applied to cancer studies, we focus on metabolic tasks that compromise cellular viability and, thus, gMCSs convert into metabolic essential genes (gMCSs of size 1) and synthetic lethals (gMCSs of size greater than 1). Importantly, gMCSs are structural properties of the reference metabolic network and, once they are obtained, we can map –omics data to identify metabolic vulnerabilities and their associated response biomarkers. Using microarray expression data, we reported a superior performance than other algorithms in the literature to predict gene essentiality, according to large-scale gene silencing data from the Project Achilles^10^.

Despite the interest in the gMCS approach since its publication^8^, further improvements are still required to make it a more practical tool in cancer research. First, there is a need for automating the application and visualization of the gMCS approach in a more intuitive and friendlier environment. Second, we need to go beyond Recon2^11^ and generate a new database of gMCSs with Human1, the most recent reference human genome-scale metabolic network^1^. Third, our previous methodology to identify cancer-specific essential genes relied on microarrays data and must be adapted to RNA-seq data, which is a more attractive and used technology for the measurement of mRNA expression.

In order to face these challenges, we present here *gMCStool*, an automated computational tool that makes use of the gMCS approach to predict metabolic vulnerabilities in cancer based on Human1 and RNA-seq data. We first show that *gMCStool* is more accurate, informative and efficient than competing approaches to predict cancer-specific essential genes. Then, a detailed illustration of *gMCStool* is presented for multiple myeloma (MM), an incurable hematological malignancy. Using different sources of RNA-seq data, which include samples from healthy donors, MM patients and cell lines, we identify metabolic liabilities of MM with *gMCStool*. *In vitro* experimental work is presented for the inhibition of CTP synthase 1 (CTPS1), a key gene involved in the pyrimidine de novo synthesis, essential for cell proliferation and viability in a group of patients with MM.

## RESULTS

*gMCStool* (https://biotecnun.unav.es/app/gmcstool) is a freely accessible web tool for the calculation of essential genes in cancer metabolism that uses genome-scale metabolic networks and RNA-seq data from human cells as input data. *gMCStool* exploits the concept of genetic Minimal Cut Set (gMCS), previously reported in Apaolaza *et al*.^8,9^. However, we introduce several major improvements in order to make gene essentiality predictions more flexible, accurate and general. The tool is organized in 5 different modules (Figure 1A): (i) ‘gMCS database’, (ii) ‘Upload RNA-seq data’, (iii) ‘Predict Essential Genes’, (iv) ‘Visualization’ and (v) ‘DepMap analysis’. A detailed illustration of the utilization of *gMCStool* can be found in the Help tab. In summary, the first 3 modules incorporate the basic functions to calculate essential genes (Figure 1B). The last two modules allow us to visualize essential genes and their companion biomarkers in the samples analyzed (Figure 1C), as well as to conduct the correlation analysis of our essentiality predictions with data from the Cancer Dependency Map (DepMap) (Figure 1D)^10,12^. Full description of these 5 modules can be found in the ‘Help’ tab of *gMCStool*. We describe below the most relevant improvements of *gMCStool* at the algorithmic level and its application for the identification of metabolic vulnerabilities in Multiple Myeloma (MM).

**Figure 1:**
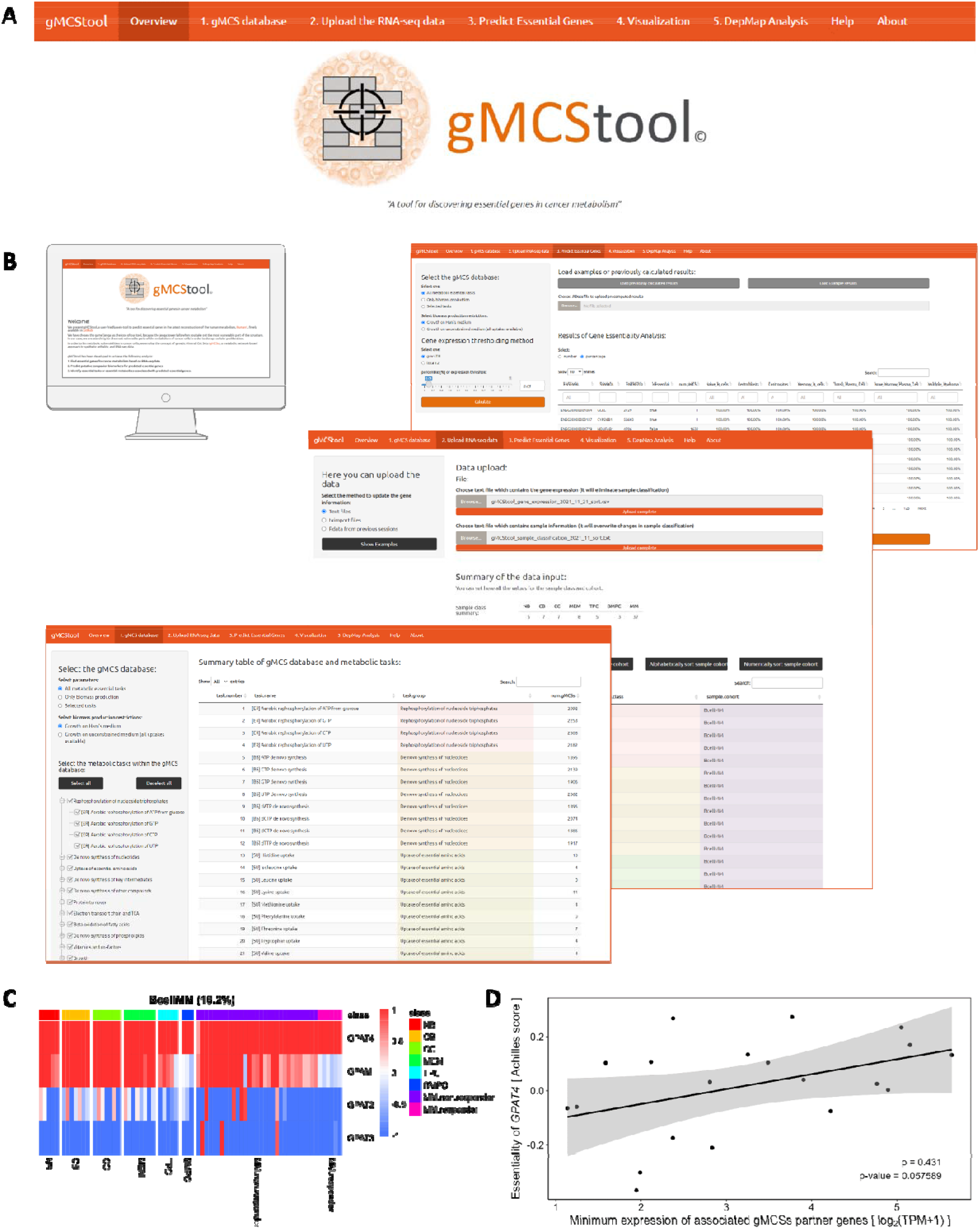
Overview of the gMCStool web application. **(A)** Snapshot of the heading of *gMCStool*, which includes the 5 main modules (‘gMCS database’, ‘Upload RNA-seq data’, ‘Predict Essential Genes’, ‘Visualization’ and ‘DepMap analysis’), ‘Help’ and ‘About’ tabs; **(B)** 3 basic modules for the calculation of essential genes in *gMCStool*. In the first module the database of gMCSs can be specified; in the second module, RNA-seq data, together with sample information, can be uploaded in different formats; in the third module, different parameters in our algorithm can be fixed and the prediction of essential genes is executed and shown in table format; **(C)** Example heatmap of gene expression data that can be obtained from the fourth module: ‘Visualization’. Expression data is shown for the predicted essential gene (*GPAT4*) and its partner genes (*GPAM, GPAT2, GPAT3*) in a specific gMCS, namely {*GPAT4, GPAM, GPAT2, GPAT3*}, for the different samples analyzed: naïve B cells (NB), centroblasts (CB), centrocytes (CC), memory B cells (MEM), tonsil plasma cells (TPC), bone marrow plasma cells (BMPC) and Multiple Myeloma (MM). It can be observed that GPAT4 is an essential gene for a subgroup of MM patients (MM responders) and for bone marrow plasma cells (BMPC) from healthy individuals. The essentiality of GPAT4 in these samples is due to the fact that its partner genes are lowly expressed; **(D)** Example dotplot that can be obtained from the fifth module: ‘DepMap analysis’, where correlation studies with DepMap are presented. In the vertical axis we have the essentiality score of *GPAT4* in DepMap for different human cell lines and in the horizontal axis the maximum expression level across partner genes: (*GPAM, GPAT2, GPAT3*). Each point represents a single cell line. In this case, only MM cell lines are shown.

### Generation of a database of gMCSs for gMCStool

Genetic Minimal Cut Sets (gMCSs) are minimal subsets of genes whose simultaneous removal directly blocks a particular metabolic task^8,9^. In cancer studies, this target metabolic task has been typically the biomass reaction, whose flux represents the proliferation rate, a key phenotype to disrupt in cancer. However, the authors of Human1 consider not only the biomass production, but also other metabolic tasks that are essential for cellular viability^1,13^ such as the production of vitamin and cofactors or activity of electron transport chain, which expands the scope of *in silico* predicted metabolic vulnerabilities. As detailed in the Methods section, we adapted our previous algorithm for the computation of gMCSs to consider the 57 metabolic tasks defined in Human1, including the biomass production (Figure 2A).

**Figure 2:**
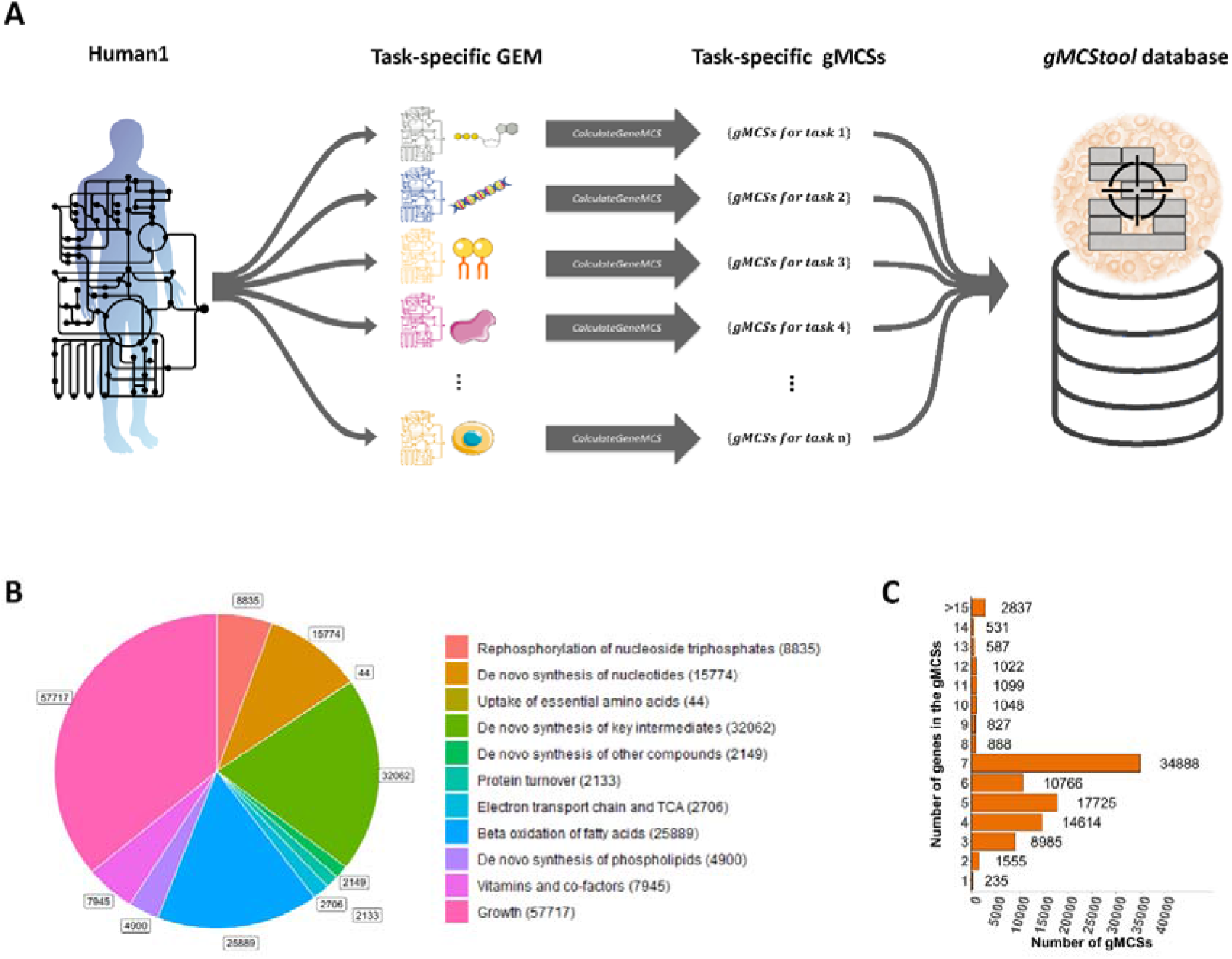
Generation of *gMCStool* database of gMCSs based on Human1. **(A)** Using Human1, the most recent reconstruction of the human metabolism, a collection of metabolic models was generated to simulate each essential metabolic task present in human cells (task-specific GEM). Then, we calculated gMCSs for each task-specific GEM (task-specific gMCSs), generating a database of synthetic lethals for human cells that are stored in *gMCStool*; **(B)** Distribution of computed gMCSs among different subgroups of metabolic tasks included in Human1; **(C)** Barplot presenting the length of gMCSs included in *gMCStool*.

As a result of our calculations, we enumerated more than 160,000 gMCSs for Human1 (see Methods section). A great part of them corresponds to biomass production (57,717); however, we also have gMCSs implied in other relevant metabolic tasks: de novo synthesis of key intermediates (32,062), beta oxidation of fatty acids (25,889) or de novo synthesis of nucleotides (15,774) (Figure 2B). Due to its simplicity or the existence of spontaneous alternative reactions, we could not find gMCSs in 5 metabolic tasks (see Supplementary Table 1). The length of computed gMCSs ranges from 1 gene to more than 15 genes, being 7 genes the most repeated solution (Figure 2C). This illustrates the high degree of metabolic flexibility of human cells. Some of them are shared across the different metabolic tasks, obtaining a total of 97,607 unique gMCSs, which overall involve 1244 metabolic genes (Supplementary Data 1). They were stored in *gMCStool* for further analysis. Supplementary Figure 1 shows the tab of *gMCStool* where the database of gMCSs, under the selected input parameters, can be downloaded. Note here that for biomass production we fixed the Ham’s growth medium, which is the one given by default in Human1 for this essential metabolic task. However, *gMCStool* provides an additional database of gMCSs for Human1 under unconstrained growth medium (all uptakes available in Human1).

### Integration of RNA-seq data into gMCStool for gene essentiality analysis

Following the concept of synthetic lethality, it is possible to predict cancer-specific essential genes by combining our database of gMCSs and gene expression data, as demonstrated in Apaolaza et al., 2017^8^. This can be done by searching for gMCSs in which all genes are lowly expressed except one of them that is highly expressed and essential for the situation under study (Figure 3A). Note here that gMCSs of size 1 directly correspond to essential genes for any cell type under the growth medium considered, in this case the Ham’s medium.

**Figure 3:**
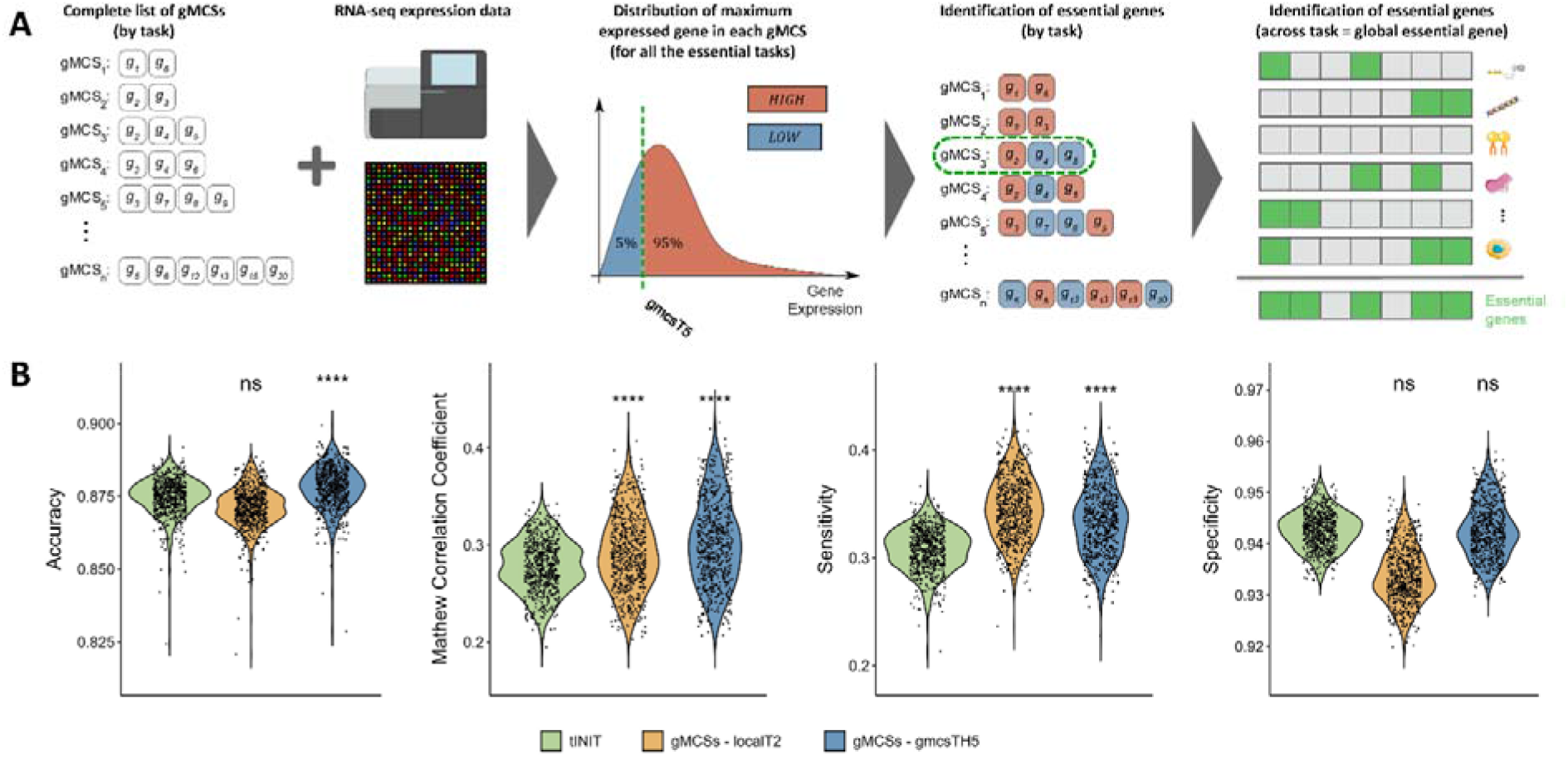
gMCS pipeline for the calculation of essential genes for Human1. (**A**) RNA-seq data is integrated with our database of gMCSs as follows. First, a potential population of highly expressed genes is obtained by extracting the maximum expressed gene of each gMCS (eliminating duplicates). Second, the X% quantile threshold (*gmcsTHX*) of this empirical probability distribution of gene expression is determined in order to discriminate between highly (in red color) and lowly (in blue color) expressed genes. For the results presented here, we fixed this threshold equal to 5% (*gmcsTH5*). Finally, in order to identify essential genes, together with gMCSs of size 1, we search for gMCSs which only contain one highly expressed gene and the rest are lowly expressed. Results are summarized in a binary matrix where columns and rows are essential genes and essential metabolic tasks. A gene is considered as essential if it is essential in at least one essential metabolic task. This process is repeated for each different sample. **(B)** Comparison of gene essentiality prediction using different methodologies and Human1 for the 621 selected cell lines with available CRISPR screening in the DepMap database ^10^. tINIT is the gold standard for CS models, and our gMCSs approach was applied with two different gene expression thresholding strategies: *gmcsTH5* and *localT2*, introduced in Richelle *et al*., 2019^14^. Note: MCC denotes Mathew Correlation Coefficient; ****/***/ns refer to the statistical significance level from an unpaired one-sided Wilcoxon test that compares the gMCS approach against the results obtained with tINIT, namely ****: p-value<=0.0001, ***: p-value<=0.001, and ns: non-significant.

In order to identify highly and lowly expressed genes in every sample using RNA-seq data, we have developed our own threshold technique for *gMCStool*, which can be applied independently to each sample (cohort-independent). Our threshold strategy exploits the fact that at least one of the genes involved in every gMCS should be highly expressed to guarantee the performance of its associated metabolic task. With this in mind, we infer for each sample a potential population of highly expressed genes by extracting the maximum expressed gene for each gMCS. Once duplicated genes were eliminated, we build an empirical probability function of the expression of highly expressed genes for each sample and fixed the X% quantile threshold of expression for them, referred as *gmcsTHX*, to alleviate possible inconsistencies and incomplete metabolic pathways in Human1 (see Methods section for details, Figure 3A). We also implemented *localT2*, a cohort-dependent methodology developed in Richelle *et al*., 2019^14^, which defines a threshold for each gene based on the observed expression distribution across the samples of the cohort. In summary, *gMCStool* incorporates these 2 thresholding approaches to categorize RNA-seq data, which is then integrated with our database of gMCSs to predict essential genes.

With the aim of assessing the prediction power of *gMCStool*, we performed a benchmark study of gene essentiality in cancer. We conducted the same analysis to the one found in Robinson *et al*., 2020^1^, based on the DepMap database, which integrates RNA-seq gene expression data^12^ and CRISPR essentiality screening experiments^10^ for a total of 621 cell lines. To avoid bias in the comparison, we used the same release of DepMap than the original analysis. As in Robinson *et al*., 2020, the genes in DepMap with Achilles score lower than -0.6 were defined as the gold-standard reference set of essential genes. We used *gMCStool* to upload RNA-seq and sample information data (Supplementary Figure 2), and to conduct gene essentiality analysis (Supplementary Figure 3). In our analysis, we considered the list of gMCSs from the 57 essential metabolic tasks (biomass production included) and predicted essential genes with both gene expression thresholding approaches: *gmcsTH5* and *localT2* (Supplementary Data 2). We compared our predicted essential genes with those resulting from DepMap (gold-standard).

As in Robinson *et al*.^1^, we calculated the accuracy, sensitivity, specificity, Matthew’s correlation coefficient (MCC). Note here that MCC is a more adequate performance metric than accuracy for cases where there is an unbalance between true positives and true negatives, as we have over 90% of non-essential genes. We also included the results presented in the publication of Human1^1^, which used *tINIT* to reconstruct 621 cell-specific GEMs and predict essential genes with single gene knockout perturbations (referred here as *tINIT*). Results can be found in Figure 3B. In the light of the MCC obtained, it can be observed that our gMCS approach overperforms tINIT with both thresholding approaches considering all metabolic tasks (unpaired one-sided Wilcoxon test p-value≤0.0001). The same result was found if we exclusively consider the essential tasks related with biomass production (Supplementary Figure 4). In addition, our *gmcsTH5* thresholding approach seems more accurate and conservative than *localT2*, which obtains the highest results in sensitivity but includes too many false negatives (unpaired one-sided Wilcoxon test p-value ≤0.0001, Supplementary Figure 4). Thus, *gMCStool* is more accurate than the state-of-the-art approach in the literature for predicting essential genes in cancer. Note here that for this gene essentiality study *gMCStool* took 36 minutes for the *gmcsTH5* approach and 31 minutes for the *localT*2 approach using a standard computer. Instead, *tINIT* required several days, as the construction of CS-models is time consuming (between 16 and 83 minutes per sample).

### Application of gMCStool to Multiple Myeloma

To illustrate the use of *gMCStool*, we performed a screening prediction of essential genes in Multiple Myeloma (MM). We used a previously generated dataset in our group that includes RNA-seq data from different B cell subpopulations from healthy individuals^15^ and bone marrow plasma cells from MM patients^16,17^. We also considered data from the MMRF-CoMMpass project, funded by the Multiple Myeloma Research Foundation (MMRF), which includes RNA-seq data for 767 MM patients at diagnosis. Finally, we obtained RNA-seq data for the 7 MM cell lines available in Cancer Cell Line Encyclopedia (CCLE)^12^.

We projected the RNA-seq expression profiles of the three datasets onto our database of gMCSs, obtaining a table which indicates the number of samples for which a gene is considered as essential in each tissue type in at least one of the essential metabolic tasks (Supplementary Data 3). We selected the MM-specific essential genes according to the following criteria: 1) to be essential in more than 10% of MM samples from our cohort; 2) to be essential in less than two samples from any healthy B cell subpopulations and less than two samples in bone marrow plasma cells, the healthy counterpart of myeloma cells; 3) to be essential in more than 5% of CoMMpass samples and 4) to be essential in one or more MM cell lines. Table 1 shows the 6 MM-specific essential genes identified in our analysis, including the percentages of samples in which the gene is considered as essential. Furthermore, we extracted the 8 gMCSs that explain the essentiality of these 6 genes in MM but only affect a few samples of B cell subpopulations (Supplementary Figures S5-S12).

**Table 1:** Specific essential genes identified in Multiple Myeloma. These genes are predicted essential in a sufficient percentage of samples of our MM cohort and MMRF-CoMMpass, but not in the samples of different B cell subpopulations obtained from healthy donors: bone marrow plasma cells (BMPC), tonsil plasma cells (TPC), memory B cells (MEM), centrocytes (CC), centroblasts (CB), naïve B cells (NB). Units are percentage of predicted samples in which a gene is essential.

| *SYMBOL* | ENSEMBL | NB<br>(n = 5) | CB<br>(n = 7) | CC<br>(n = 7) | MEM<br>(n = 8) | TPC<br>(n = 5) | BMPC<br>(n = 3) | MM<br>(n = 37) | CoMMpass<br>(n = 767) | CCLE<br>(n = 7) |
| --- | --- | --- | --- | --- | --- | --- | --- | --- | --- | --- |
| *CTPS1* | ENSG00000171793 | 0% | 0% | 0% | 0% | 0% | 0% | 11% | 71% | 86% |
| *DHFR* | ENSG00000228716 | 0% | 0% | 0% | 0% | 20% | 0% | 11% | 57% | 86% |
| *LCAT* | ENSG00000213398 | 0% | 29% | 0% | 0% | 0% | 0% | 11% | 7% | 14% |
| *NMNAT3* | ENSG00000163864 | 0% | 0% | 0% | 0% | 0% | 0% | 16% | 28% | 14% |
| *PC* | ENSG00000173599 | 0% | 0% | 0% | 0% | 0% | 0% | 32% | 17% | 86% |
| *UAP1* | ENSG00000117143 | 0% | 0% | 0% | 0% | 0% | 0% | 16% | 14% | 29% |

From the results shown in Table 1, we focused on CTPS1 for further analysis. Figure 4A shows the gMCS that involves *CTPS1* and *CTPS2* and suggests the essentiality of CTPS1 in a subgroup of MM samples but not in the healthy tissues from different B cell subpopulations. Similarly, it can be observed the patients from CoMMpass that could be responders and non-responders to CTPS1 inhibition based on the expression of CTPS2. The same result can be observed for the 7 MM cell lines considered. In addition, Figure 4B shows that the expression of *CTPS2* decreases for the sub-group of MM samples that potentially could respond to CTPS1 inhibition (MM responders). Note here that we used *gMCStool* to automatically generate these figures (Supplementary Figure 13).

**Figure 4:**
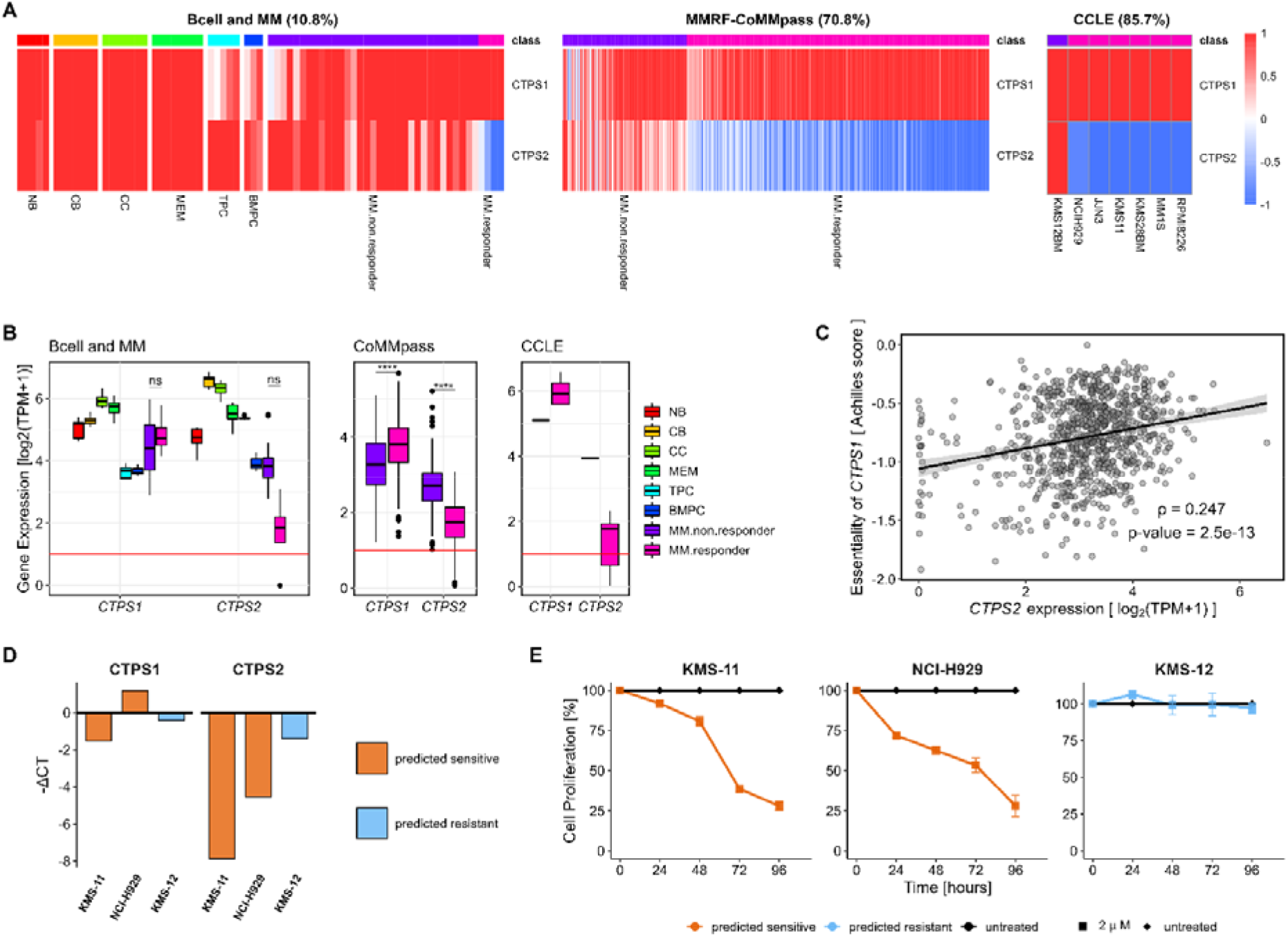
Prediction of essentiality of CTPS1 in MM with *gMCStool*. **(A)** Selected gMCS: {*CTPS1*, *CTPS2*} and gene expression in our cohort of patients, MMRF-CoMMpass and selected MM Cell Lines; **(B)** Boxplot of expression of the *CTPS1* and *CTPS2* genes in the B cell differentiation and MM samples; **(C)** Correlation between the essentiality of *CTPS1* (CRISPR knockout screen data, DepMap) and the expression of *CTPS2* in log2(TPM+1). **(D)** qPCR-RT of *CTPS1* and *CTPS2* expression in the selected cell lines. Data are referred to GUS gene. Full experimental results are found in Supplementary Table 3. **(E)** Proliferation of H929, KMS11 and KMS12 cell lines treated with CTPS1 inhibitor. The proliferation percentage refers to non-treated cells (black line). Data represent mean◻±◻standard deviation of at least three experiments. Note: ****/ns refer to the statistical significance level from an unpaired one-sided Wilcoxon test, namely ****: p-value<=0.0001 and ns: non-significant.

At the center of our hypothesis above is that CTPS1 and CTPS2 are synthetic lethal and, thus, the essentiality of CTPS1 depends on CTPS2, namely when CTPS2 is lowly expressed, CTPS1 becomes essential. We assessed this hypothesis with DepMap data available in *gMCStool*, finding a positive and significant correlation between the *Achilles* score of CTPS1 and the expression of *CTPS2* (rho = 0.247, p-val = 2.5e-13, Supplementary Figure 14), as detailed in Figure 4C. This provides further support to our hypothesis of synthetic lethality of CTPS1 and CTPS2.

Furthermore, *gMCStool* provides the associated metabolic task for each gMCS. In this particular case, the inhibition of CTPS1 and CTPS2 blocks several tasks: CTP (*cytidine triphosphate*) de novo synthesis, dCTP (*deoxycytidine triphosphate*) de novo synthesis and biomass production, which turns out to disrupt the production of CTP, dCMP (*deoxycytidine monophosphate*), DNA and RNA. Based on this information, we performed an *in silico simulation for* the addition of cytidine to the growth medium, showing a rescue of proliferation after the inhibition of *CTPS1* and *CTPS2*, which illustrates how *gMCStool* provides further evidence for predicted synthetic lethals (Supplementary Table 2).

Finally, we carried out *in vitro* experimental validation of the essentiality of *CTPS1* in three MM cell lines. For two of them, H929 and KMS11, *gMCStool* had predicted their sensitivity to CTPS1 inhibition due to their low expression of *CTPS2*, while for one of them, KMS12, *gMCStool* had predicted its resistance to *CTPS1* inhibition due to its high expression of *CTPS2* (Figure 4D, Supplementary Table 3). Specifically, we synthetized a CTPS1 inhibitor, previously developed by Rao and colleagues^19^, and assessed its effect in the proliferation of the three cell lines mentioned above. As a result, H929 and KMS11 reduced their proliferation more than 50% after 4 days of culture and no alterations were obtained for KMS12 (Figure 4E). These positives results reinforce the predictive power of *gMCStool* and open new research avenues to treat MM.

## DISCUSSION

In this work, we present *gMCStool*, a computational tool for the prediction of metabolic vulnerabilities in cancer based on gMCSs, a network-based approach to synthetic lethality, and RNA-seq data. *gMCStool* incorporates technical improvements in our previously developed algorithms^8,9^ and addresses the need of efficiently automating the application and visualization of the gMCS approach in a simple and intuitive environment.

Importantly, *gMCStool* stores more than 160,000 gMCSs that block at least one essential metabolic task of Human1, the most recent genome-scale metabolic network of human cells. The computation of this database of gMCSs substantially simplifies the process of identifying cancer-specific essential genes, which can be now extracted by correctly mapping gene expression data onto them. This strategy makes gene essentiality analysis more accessible and natural to researchers less familiar with the field of constraint-based modeling.

In addition, *gMCStool* allows us to perform gene essentiality analysis more efficiently than other algorithms in the literature, as we avoid the step of constructing context-specific metabolic models, which makes use of time-consuming optimization techniques. For example, *gMCStool* required 36 minutes to calculate the essential genes for all the 621 cancer cell lines available in DepMap, whereas tINIT needs between 30-60 minutes to reconstruct the metabolic model of one single cell line using RNA-seq data. Thus, *gMCStool* substantially reduces the computational requirements to conduct gene essentiality analysis in cancer. Specifically, we could analyze the samples in the Results section, which add up to more than 1400 samples, considering DepMap and different MM cohorts, in less than 80 minutes with a standard computer. Overall, *gMCStool* constitutes an effective and friendly online tool to search for metabolic vulnerabilities in cancer research.

In addition to make *gMCStool* a practical tool for researchers in cancer metabolism, we extended our previously developed algorithms in order to: 1) consider that Human1 involves different essential metabolic tasks beyond biomass production, typical target in network-based gene essentiality analysis; 2) predict essential genes based on RNA-seq data, namely by proposing a novel approach to discriminate between highly and lowly expressed genes (*gmcsTHX*). These advances allowed us to compare the accuracy of *gMCStool* with tINIT^1,13,20^, the approach developed by the authors of Human1 in order to build CS metabolic models of cancer cells and conduct gene essentiality analysis. Importantly, this study shows that *gMCStool* is significantly more accurate that tINIT when compared with gene essentiality screens available in DepMap.

Another advantage of *gMCStool* is its visualization capabilities. *gMCStool* exploits the fact that essential genes are derived from specific gMCSs, where one gene is highly expressed and the rest genes are lowly expressed. Thus, genes involved in gMCSs do not only allow us to predict essential genes, but also response biomarkers for their inhibition. This idea underlies the different plots that can be extracted from *gMCStool*, which facilitates the interpretation of our computational predictions. In this direction, *gMCStool* also outputs essential tasks and metabolite biosynthesis that are disrupted by gMCSs, which are particularly informative to construct testable hypotheses about the mechanism behind predicted synthetic lethals. Note here that this might be of interest due to the fact that the composition of the human biomass is typically defined as universal for all cells, but some authors think that this might be an incorrect assumption^21–25^, suggesting that it could be context-specific and some metabolites are more relevant than others for different tumors.

To exemplify the use of *gMCStool* in personalized medicine, we performed gene essentiality analysis in the different B-cell differentiation subpopulations and MM samples from several cohorts, aiming to identify candidates that maximizes the number of MM samples affected but, at the same time, minimizes the toxicity of the treatment, which is simulated here as the number of healthy tissue samples affected by such target. Despite the large number of gMCSs stored in gMCStool, we only spot 6 metabolic enzymes whose inhibition is selectively toxic for MM cells, which illustrates the difficulty of identifying cancer-specific metabolic processes. Among them, we focused on *CTPS1*. *In vitro* experiments confirmed the predictions of *gMCStool* in 3 MM cell lines: 2 positives for *CTPS1* inhibition and 1 negative for *CTPS1* inhibition, according to the expression of *CTPS2*. These results support the synthetic lethality of *CTPS1* and *CTPS2* and open new research area to treat MM.

Finally, despite the advance of *gMCStool* over existing tools, there is still a long way to achieve the desired performance in predicting cancer-specific essential genes using genome-scale metabolic networks. *gMCStool* reached a maximum of 35% of sensitivity and F1 score using DepMap as gold-standard. The extension and update of existing reference genome-scale metabolic networks is obviously a critical task to improve further accuracy metrics. In this respect, Human1 has established an open and active community that provides a successful and integrative response to that need. In addition, the definition of cancer-specific metabolic tasks, beyond common essential metabolic tasks, such as biomass production^26^, could open new avenues to identify essential genes and increase sensitivity. We recently illustrated the relevance of polyamines in different hematological tumors, and these metabolites are not typically considered in standard biomass reactions^27^. The development of systematic methods to extract these essential metabolites in different contexts constitutes a challenging issue. Finally, the integration of metabolic and regulatory networks is a difficult but relevant task that has a great impact in predicting essential genes, particularly when the proxy for activity is gene expression data that could be modified due to compensatory regulatory pathways. Making progress in all these challenges will help not only *gMCStool*, but all methods in the literature to predict more accurately essential genes and response biomarkers in order to address unmet clinical needs and reach our goal to provide patients more personalized treatments.

## METHODS

### Calculation of genetic Minimal Cut Sets (gMCSs) in Human 1

The reference metabolic network Human1 (version 1.4.0)^1^ was obtained from https://github.com/SysBioChalmers/Human-GEM. Human1 involves 13,416 reactions, 8,458 metabolites and 3,628 genes. Importantly, Human1 defines 56 essential metabolic tasks for any human cell in addition to the biomass production, typically included in other genome-scale metabolic reconstructions^11,28,29^. Each of them defines a list of output metabolites that must be derived from a list of input metabolites and, if necessary, an artificial equation that is required to support this transformation. Lower and upper bounds are also fixed for a specific subset of reactions involved in each metabolic task.

With this information, a different metabolic model for each essential task can be built and used to assess the effect of genetic perturbations via linear programming, namely by assuming the mass balance condition and thermodynamic constraints^30^. Essential genes correspond to those single gene knockouts leading to an infeasible linear programming (at least one of the required lower/upper bounds in the metabolic task is violated). However, this strategy is not efficient to calculate higher order essential gene knockout perturbations (synthetic lethals), due to the combinatorial nature of the problem. The gMCS approach is suitable for this task^9^.

For the calculation of gMCSs, we used the *calculateGeneMCS* function available in the COBRA Toolbox^31^, available at https://github.com/opencobra/cobratoolbox/. This function requires a metabolic model and a single target reaction to be blocked as input data. We constructed this input data for each metabolic task following its associated information about inputs, outputs, artificial equations and lower bounds. A detailed illustration as to how the target reaction was derived for each metabolic task can be found in Supplementary Note 1.

Note here the adaptation of Human1 to the *calculateGeneMCS* function from COBRA leads to slightly different metabolic models to the ones originally built Human1. However, we checked for each metabolic task that the essential genes in our metabolic models, obtained with the *singleGeneDeletion* function available in COBRA, and the original models in Human1, derived from the *checkTasksGenes* function from RAVEN^32^, were the same.

In order to reduce the computational cost of calculating gMCSs, the resulting metabolic models for each essential task were simplified using the *fastFVA* function from COBRA (without any requirement in the objective function). Finally, all calculated gMCSs were checked using the *checkTasks* function available in RAVEN using the original metabolic models in Human1. This task was performed in order to remove possible false positive gMCSs that could arise due to the time limit fixed in the Mixed-Integer Linear Programming (MILP) solver used in the *calculateGeneMCS* function ^9^. These results were computed with Intel(R) Xeon(R) Silver 4110 CPU @ 2.10GHz processors, limiting to 8 cores and 8 GB of RAM. A time limit of 120 seconds was set for each solution derived from the function *CalculateGeneMCS*.

### Gene categorization based on RNA-seq data

In our computational approach, we denote a gene as essential in a particular sample when it is the only gene expressed in at least one gMCS. Thus, in order to identify essential genes for each sample, we need to systematically decide which genes are highly (ON) or lowly (OFF) expressed. To that end, we developed our own methodology using RNA-seq expression data.

Our approach exploits our database of gMCSs by assuming that all of them should involve at least one highly expressed gene to guarantee that their associated metabolic tasks are feasible. In particular, we assume that the gene with the highest expression in each gMCS is the one that should be highly expressed. Thus, we extracted the maximum expression level in every gMCS and every essential task and generated an empirical distribution of highly expressed genes for each sample (**X_H_**). Note here that repetitions are not taken into account and each gene contributes exactly one value to the distribution. We considered as highly expressed those genes expressed above the Xth percentile, referred here as *gmcsTHX*, and lowly expressed otherwise. We fixed the threshold of expression at 5th percentile (*gmcsTH5*) to alleviate possible inconsistencies and incomplete metabolic pathways in Human1.

In addition, we used the *localT2* methodology^14^. To put it briefly, a gene is considered as MAYBE ON in a specific sample whenever its expression level is greater than its mean expression level across the samples of the cohort, MAYBE OFF otherwise. Additionally, two global expression thresholds are applied in *localT2*: genes whose expression is below the 25th percentile of the distribution of expression for all samples and genes are considered as OFF, whereas those above the 75th percentile of the same distribution are considered as ON. This global expression threshold dominates the categorization obtained from the relative expression threshold based on the mean expression value. Note here that our *gmcsTH5* approach is independently applied to each sample, *i.e*. we identified *gmcsTH5* for each sample and defined the subset of highly and lowly expressed genes as those having an expression value higher and lower than *gmcsTH5*, respectively. Instead, *localT2* is dependent on the cohort to categorize genes, as they consider all samples to establish global and relative thresholds. Both thresholding methods are available in *gMCStool*. They were applied to categorize the 1244 genes that participate in all the calculated gMCSs in the different cohorts of patients and cell lines considered in the Results section. A sensitivity analysis was performed in Supplementary Figure 15, which shows the same analysis presented in Figure 2 for several percentiles of *gmcsTHX* (*gmcsTH0, gmcsTH1, gmcsTH5, gmcsTH10*), finding robust results for lower expression thresholds. Moreover, we also tested the implication of normalization of TPM for *localT2* and the effect of the gene-set selection for the 25th-75th threshold of *localT2*.

### Gene essentiality analysis using gMCSs and gene expression data

Once all gMCS have been calculated and the gene expression values are classified as highly or lowly expressed, gene essentiality analysis can be carried out. In particular, we search for gMCSs which have exclusively one gene defined as highly expressed, and the rest of genes lowly expressed. For each gene and each task, we identify gMCSs that contain that given gene as highly expressed and the rest as lowly expressed. If there is at least one gMCS that fulfills this condition, the gene is considered as essential. *gMCStool* can create a list of essential genes and associated gMCSs for each essential metabolic task in Human1. Note here that gMCSs that comprise only one gene are essential genes in all human cells.

### Multiple Myeloma case study

RNA-seq data from our group consist of 37 MM patients (GSE151063)^16,17^ and 35 samples from different B cell subpopulation (GSE114816)^15^. Sample 57802 was removed from the study for being detected as an outlier in a PCA analysis. RNA-seq from the MMRF-CoMMpass has 767 samples from MM patients at diagnosis time, available in IA18 release. RNA-seq from 7 MM cell lines was obtained from DepMap, release 21Q1^12^. These cell lines are NCIH929, JJN3, KMS11, KMS12BM, KMS28BM, MM1S and RPMI8226. RNA-seq data was downloaded in TPM and profiled using both threshold methodologies: *gmcsTH5* and *localT2*.

### Metabolite essentiality prediction

In order to identify essential metabolites whose production is disrupted through gMCSs, we only focused on the biomass production task, due to its underlying complexity. The rest of essential metabolic tasks are well-defined and highly specific, depending only on a reduced number of metabolites.

To that end, we first defined a list of metabolites directly related to the biomass production. As in the biomass reaction of Human1 there are several metabolite pools, we also included their precursors in such target list. Then, we created a sink exchange reaction for each target metabolite and assessed their production through an FBA analysis^30^ after the removal of genes involved in each gMCS. We extracted for each gMCS the essential metabolites whose production is disrupted (maximal production is zero).

### Q-PCR

The expression of *CTPS1* and *CTPS2* were analyzed by Q-PCR in MM KMS-11, KMS-12 and NCI-H929 cell lines. RNA was extracted with TRIzol Reagent (Invitrogen) according to the manufacturer’s instructions. First, cDNA was synthesized from 1 μg of total RNA using the PrimeScript RT reagent kit (Perfect Real Time) (Cat No RR037A, TaKaRa) following the manufacturer’s instructions. The quality of cDNA was checked by a multiplex PCR that amplifies *PBGD*, *ABL*, *BCR* and *β2-MG* genes. Q-PCR was performed in a QuantStudio 5 Real-Time PCR System (Applied Biosystems), using 20 ng of cDNA in 2 μL, 1 μL of each specific primer at 10 μM (CTPS1 F: TTATTGAGGCCTTCCGTCAG; CTPS1 R: GGGAAAGCCCAAGTCCTCTA; CTPS2 F: GCTGTCCAGGAGTGGGTTAT; CTPS2 R: CGCCTTAAACTGGAATTGTCT), 5 μL of SYBR Green PCR Master Mix 2X (Cat No 4334973, Applied Biosystems) in 10 μL reaction volume. The following program conditions were applied for Q-PCR running: 50 °C for 2 min, 95 °C for 60 s following by 45 cycles at 95 °C for 15 s and 60 °C for 60 s; melting program, one cycle at 95 °C for 15 s, 40 °C for 60 s and 95 °C for 15 s. The relative expression of each gene was quantified by the Log 2(-ΔΔCt) method using the gene *GUS* as an endogenous control.

### Cell culture

KMS-11, KMS-12 and NCI-H929 cell lines were maintained in culture in RPMI1640 medium (Gibco, Grand Island, NY) supplemented with 10% fetal bovine serum (Gibco, Grand Island, NY) and penicillin/streptomycin (BioWhitaker, Walkersvill, MD) at 37◻°C in a humid atmosphere containing 5% CO2. Cell lines were obtained from the DSMZ or the American Type Culture Collection (ATCC). All cell lines were authenticated by performing a short tandem repeat allele profile and were tested for mycoplasma (MycoAlert Sample Kit, Cambrex), obtaining no positive results.

### Small molecule synthesis of CTPS1 inhibitor

Synthesis of compound 1 from Rao *et al*.^19^ was performed by Wuxi Apptec and consisted of three steps. First, to a solution of EDCI (172.07 mg, 897.57 umol, 1.5 eq) and DMAP (73.10 mg, 598.38 umol, 1 eq) in DCM (5 mL) were added 3-methoxyaniline (110.54 mg, 897.57 umol, 100.49 uL, 1.5 eq). This solution was then added to 3-nitrobenzoic acid (0.1 g, 598.38 umol, 1 eq) and the solution was stirred at 25°C for 3 hours. TLC (Petroleum ether: Ethyl acetate=0:1) indicated the reaction was completed and one new spot formed. The reaction was clean according to TLC. The reaction mixture was quenched by addition water 5 mL. The organic layer was separated and washed with 1M aqueous HCl 5 mL, dried over Na_2_SO_4_, filtered and concentrated under reduced pressure to give a residue. Compound N-(3-methoxyphenyl)-3-nitro-benzamide (150 mg, 550.95 umol, 92.07% yield) was obtained as a yellow solid. Second, to a solution of N-(3-methoxyphenyl)-3-nitro-benzamide (150 mg, 550.95 umol, 1 eq) in THF (8 mL) was added Pd/C (50 mg, 5% purity,) under N_2_. The suspension was degassed under vacuum and purged with H_2_ several times. The mixture was stirred under H2 (15 psi) at 25°C for 2 hours. TLC (Petroleum ether/Ethyl acetate=3:1) showed the starting material was consumed completely. The reaction mixture was filtered and the filtrate was concentrated. Compound 3-amino-N-(3-methoxyphenyl) benzamide (100 mg, 412.76 umol, 74.92% yield) was obtained as a white solid. Finally, to a solution of 3-amino-N-(3-methoxyphenyl)benzamide (100 mg, 412.76 umol, 1 eq) in DCM (5 mL) was added DIEA (106.69 mg, 825.52 umol, 143.79 uL, 2 eq) and 2-chloroacetyl chloride (46.62 mg, 412.76 umol, 32.83 uL, 1 eq). The mixture was stirred at 0 °C for 1 hr. LC-MS showed the reaction was completed and one main peak with desired m/z was detected. The reaction mixture was quenched by addition H_2_O 10 mL, and extracted with DCM 10 mL (5 mL * 2). The combined organic layers were washed with brine 10 mL, dried over Na_2_SO_4_, filtered and concentrated under reduced pressure to give a residue, which was washed by MeCN (10 mL), then filtered to collect the solid. Compound 3-[(2-chloroacetyl)amino]-N-(3-methoxyphenyl) benzamide (50.49 mg, 155.04 umol, 37.56% yield, 97.879% purity) was obtained as an off-white solid. ESI-MS m/z: calcd for C_16_H_15_ClN_2_O_3_ 318.08, m/z found 319.1 [M+H]^+^. ^1^H-NMR (DMSO, 400MHz): δ ppm 10.50 (br s, 1H), 10.25 (br s, 1H), 8.09 (br s, 1H), 7.82 (br d, *J* = 8.4 Hz, 1H), 7.66 (br d, *J* = 7.2 Hz, 1H), 7.54 – 7.42 (m, 2H), 7.36 (br d, *J* = 8.0 Hz, 1H), 7.30 – 7.21 (m, 1H), 6.69 (br d, *J* = 8.0 Hz, 1H), 4.28 (s, 2H), 3.75 (s, 3H).

### CTPS1 inhibitor treatment and cell proliferation assay

KMS-11, KMS-12 and NCI-H929 cell lines were treated with 2μM of the CTPS1 inhibitor for 24, 48, 72 and 96 hours. After the indicated times of treatment, cell proliferation was analyzed using the CellTiter 96 Aqueous One Solution Cell Proliferation Assay (Promega, Madison, W) following the manufacturer’s instructions. First, the average of the absorbance from the control wells was subtracted from all other absorbance values. Data were calculated as the percentage of total absorbance of treated cells/absorbance of non-treated cells.

### Implementation and availability

*gMCStool* has been developed using R^33^ and Shiny^34^. *gMCStool* is hosted using the Amazon Web Services cloud environment service and it can be publicly accessed in: https://biotecnun.unav.es/app/gmcstool. A full tutorial and example for our own cohort of samples corresponding to B cell differentiation and MM can be found in the ‘Help’ tab of *gMCStool*.

### Code availability

The code for *gMCStool* is available in https://github.com/lvalcarcel/gMCStool. Using this tool, it is possible to generate all the results presented in this article. Gene expression data should be a matrix which have genes (in ENSEMBL annotation) for rows and samples for columns.

### Accession numbers and datasets

The authors declare that all data supporting the findings of this study are available within the article, in the supplementary material or in other studies.

Referenced accession: B-cell and MM RNA-seq data was obtained from GEO under accession codes GSE151063^16,17^ and GSE114816^15^. MMRF-CoMMpass data were generated as part of the Multiple Myeloma Research Foundation Personalized Medicine Initiatives (https://research.themmrf.org and www.themmrf.org). All cancer cell lines are public and accessible in www.depmap.org^10,12^ (release 21Q2).

## ACKNOWLEDGEMENTS

This work was supported by the Minister of Economy and Competitiveness of Spain [PID2019-110344RB-I00], PIBA Programme of the Basque Government [PIBA_2020_01_0055], Elkartek programme of the Basque Government [KK-2020/00008], Fundación Ramon Areces [PREMMAM] to F.J.P.; Instituto de Salud Carlos III (ISCIII) [PI16/02024, PI17/00701, PI19/01352, PI20/01306], CIBERONC (Co-financed with European Union FEDER funds) [CB16/12/00489], ERANET program ERAPerMed [MEET-AML], MINECO Explora [RTHALMY], Cancer Research UK and AECC under the Accelerator Award Programme [C355/A26819]. to F.P. and Instituto de Salud Carlos III (ISCIII) [FI17/00297] to L.V.V.

## AUTHORSHIP CONTRIBUTIONS

X.A, F.P., and F.J.P. conceived this study. L.V.V., I.A. and F.J.P. developed the conceptual approach and L.V.V. performed the computational analysis. E.S.J.-E., R.O., A.V., L.G., A.P-L and X.A. performed the experiments. F.P., X.A, E.S.J.-E. and J.S-M. analyzed and provided access to multiple myeloma data. All authors wrote, read, and approved the manuscript.

## DISCLOSURE OF CONFLICTS OF INTEREST

The authors declare no competing financial interests.

## Notes

### Competing Interest Statement

The authors have declared no competing interest.

